# Infections patterns and fitness effects of *Rickettsia* and *Sodalis* symbionts in the green lacewing *Chrysoperla carnea*

**DOI:** 10.1101/327130

**Authors:** Rebekka Sontowski, Michael Gerth, Sandy Richter, Axel Gruppe, Martin Schlegel, Nicole van Dam, Christoph Bleidorn

**Affiliations:** German Centre for Integrative Biodiversity Research (iDiv) Halle-Jena-Leipzig, Germany; Institute of Biodiversity, Friedrich-Schiller-University, Jena, Germany; Institute of Integrative Biology, University of Liverpool, Liverpool, UK; Sobell Department of Motor Neuroscience & Movement Disorders, UCL Institute of Neurobiology, University College London, Queen Square, London WC1N 3BG, UK; Institute of Biology, Molecular Evolution and Systematics of Animals, University of Leipzig, Leipzig, Germany; Chair of Zoology – Entomology group, Technical University of Munich, Freising, Germany; Animal Evolution and Biodiversity, University of Göttingen, Göttingen, Germany

**Keywords:** biological pest control, co-infection, endosymbiont, Neuroptera, Rickettsiales, symbiosis

## Abstract

Endosymbionts are wide-spread among insects and can play an essential role in host ecology. The common green lacewing (*Chrysoperla carnea* s. str.) is a neuropteran insect species which is widely used as a biological pest control. We screened for endosymbionts in natural and laboratory populations of the green lacewing using diagnostic PCR amplicons. We found the endosymbiont *Rickettsia* to be very common in all screened populations, whereas a so far uncharacterized *Sodalis* strain was solely found in laboratory populations. The new *Sodalis* strain was characterized using a whole genome shotgun approach. Its draft genome revealed an approximate genome size of 4.3 Mbp and the presence of 5213 coding sequences. Phylogenomic analyses indicated that this bacterium is the sister taxon of *S. praecaptivus*. In an experimental approach, we found a negative impact of *Sodalis* on the reproduction success of the green lacewing. Co-infections with *Rickettsia* and *Sodalis* caused an even higher decrease of reproductive success than single *Sodalis* infections. In contrast, no significant fitness differences were found in *Rickettsia* infected green lacewings compared to uninfected lacewings. The *Rickettsia/Sodalis/Ch. carnea* system presents a promising model to study evolutionary endosymbiont-host interactions in Neuroptera and endosymbiont-endosymbiont interactions in general. The economic and ecological importance of green lacewings in biological pest control warrants a more profound understanding of its biology, which might be strongly influenced by symbionts.

## Introduction

With about 6000 species Neuroptera represent a rather small group of insects [1]. One well-known representative of the Neuroptera is the common green lacewing *Chrysoperla carnea*. Originally assumed to represent a single species [2], *Ch. carnea* s. lat. was shown to be a species complex, which are morphologically difficult to distinguish [3]. The adults of these species feed on honey dew and pollen while the larvae are predators of a broad range of insects, e.g. aphids, mealy bugs and other soft-bodied species [4,5] Fittingly, lacewing larvae are efficient biological pest control agents in the field, greenhouses and orchards [6,7]. Biological pest control has received much attention through increasing insecticide resistance of several pests and legislations that aim to reduce usage of synthetic chemical pesticides. Green lacewing larvae possess a high resistance against many widely used pesticides and because of their usefulness in pest control, lacewings are mass-reared and marketed commercially [7,8].

Endosymbionts are wide-spread among insects and can play an essential role in host ecology. Obligate endosymbionts are essential for their insect hosts to survive, e.g. by providing essential nutrients [9,10]. Facultative associates are not essential for their host, but impact host fitness through various induced interactions, e.g. reproductive manipulation and color modifications [11,12] One of the most common endosymbionts in insects is *Rickettsia* sp. (α-Proteobacteria) with an estimated occurrence in one quarter of all terrestrial arthropod species [13]. *Rickettsia* sp. infects vertebrates, arthropods and plants [14,15]. Several lineages occur in vertebrates, e.g. as human pathogens, and are transmitted by arthropods [14]. However, the majority of lineages are exclusively found in arthropods [16]. Some of them are able to manipulate host reproduction by causing male killing in ladybird beetles (Coccinellidae) and jewel beetles (Buprestidae), or parthenogenesis in eulophid wasps [17–20]. *Rickettisa* sp. is abundant in natural and laboratory insect populations, establishes itself rapidly in populations, and remains stable at high frequencies [21,22].

Endosymbionts in Neuroptera have so far been largely neglected. Two recent studies have described male-killing *Spiroplasma* in the green lacewing *Mallada desjardinsi* [23,24]. *Rickettsia* infections were first described in randomly sampled arthropod host screens that included Neuroptera [16,25]. Recently, a Neuroptera-specific *Rickettsia* screening showed that approximately 40% of the tested Neuroptera species were infected, including *Ch. carnea* s. str. [26] which was infected by strains of the *R. bellii* clade, commonly found in arthropods [16,26]. While screening *Ch. carnea* s. str. for endosymbionts, we also found infections with *Sodalis* sp., a facultative endosymbiont belonging to the γ-proteobacteria [27]. *Sodalis* was first identified in tsetse flies and later detected in different insect groups such as weevils, stinkbugs, louse flies and lice [28–32]. The prevalence of *Sodalis* infections can vary greatly [33]. The reported host-*Sodalis* endosymbiont interactions are highly diverse, ranging from facultative to obligate [34]. They are able to increase trypanosome infections in tsetse flies (Glossinidae), participate in the cuticle synthesis of weevils and modify host phenotypes [35,31]. Even more complex, a low prevalence of *Sodalis* in weevils produces a host killing phenotype, whereas a high prevalence leads to a persistent and beneficial infection in the hosts [36].

Based on the successful application of green lacewings as biocontrol, they are commercially mass-reared. However, it is still unclear how these endosymbionts are distributed on species and population levels and which role they play in those hosts. In the present study, we screened several natural populations of *Ch. carnea* s. str. for *Rickettsia* and *Sodalis* infections. Given their effective application as biological pest control, *Ch. carnea* is commercially available. We therefore also tested if infection patterns in natural and laboratory populations are similar. By doing so, we discovered that the commercial lines also contained a *Sodalis* species. To characterize the *Sodalis* symbiont in *Ch. carnea* s. str., we assembled a draft genome from Illumina short reads for subsequent phylogenomic analysis. Moreover, host-endosymbiont interactions were investigated by generating *Ch. carnea str.* lines that carried *Rickettsia* or *Sodalis*, or both, as well as endosymbiont-free lines. Based on these cultures, we examined the rate of vertical transmission of the symbionts and their potential impact on host reproduction. We demonstrate that the *Ch. carnea str./Rickettsia/Sodalis* system represents a promising model for the evolution of insect-endosymbiont interactions in general.

## Material and methods

### Population level endosymbiont screening in natural and laboratory *Ch. carnea* s. str

To compare the occurrence of endosymbionts under natural conditions to those from commercially reared laboratory populations, we obtained *Ch. carnea* s. str. individuals from three different companies (N=64 in total) and sampled eight natural populations (N=84 in total). The supplying companies were Sautter & Stepper GmbH (Ammerbuch-Altingen, Germany, 26 larvae), Biobest (Westerlo, Belgium, 18 larvae) and Katz Biotech AG (Baruth/Mark, Germany, 20 larvae). Additional adult lacewings were collected from various locations in Germany and Austria between 2010 and 2015 (Table S1). DNA was extracted from whole insects using the NucleoSpin Tissue Kit (MARCHERY-NAGEL, Düren, Germany). To identify a *Rickettsia* and/or *Sodalis* infections, a PCR screening with species-specific *16S* rRNA primer and PCR programs was performed (Table S2). Amplicons were counted as positive evidence for *Rickettsia* or *Sodalis* infection. To exclude false negatives, the PCR product was diluted to 1:10, 1:100, and 1:1000. These dilutions were subjected to PCR again. When no bands were visible for any of the PCRs, the sample was counted as not infected.

### Molecular characterization of endosymbionts

*Rickettsia* endosymbionts of the investigated green lacewings were already phylogenetically characterized in Gerth et al. [26]. For the *Sodalis* endosymbiont, we used a whole genome shotgun approach to generate a draft genome for phylogenomic analyses. For this purpose, a double-indexed Illumina library from *Sodalis* infected second instar green lacewing larvae DNA was constructed as detailed in Meyer and Kircher [37] and Kircher et al. [38]. The insect sample was obtained from the company Sautter and Stepper GmbH (Ammerbuch-Altingen, Germany). The library was sequenced as 140-bp paired-end run on an Illumina HighSeq 2500 (Illumina, San Diego, CA) at the Max Planck Institute for Evolutionary Anthropology (Leipzig, Germaany). Base calling was performed with freeIbis [39], adapters were trimmed and reads with more than five bases below a quality threshold of 15 were discarded. A preliminary meta-assembly was created using IDBA-UD [40], with k-mers 21–81 in steps of ten. Assembled contigs were blasted with BLASTN against NCBI GenBank. All recovered *Sodalis* contigs were used as reference for subsequent mapping using NextGenMap 0.4.12 [41] to retrieve all putative *Sodalis* reads. The coverage of all *Sodalis* contigs was evaluated with qualimap 2.2.1 [42] and retrieved reads were newly assembled with SPAdes 3.1.1 [43], an assembler optimized for bacterial and archaeal genomes, to generate a *Sodalis* draft genome. Raw reads and assembly have been submitted to NCBI Genbank under the accessions TO BE ADDED.

For phylogenomic analysis, genome assemblies of representative *Sodalis* lineages were downloaded from NCBI (*Candidatus Sodalis* sp. *Socistrobi* (GCA_900143145); *S. glossinidius* (GCA_000010085); *Sodalis* symbiont of *Philaenus spumarius* (GCA_000647915); *S. pierantonius* (GCA_000647915); *S. praecaptivus* (GCA_000517425); *Sodalis* symbiont of *Proechinophthirus fluctus* (GCA_001602625); *Sodalis* sp. TME1 (GCA_001879235)), as well as five selected outgroups (*Eschericha coli* K12 (NC_000913); *Pectobacterium carotovorum* (NC_012917); *Photorhabdus luminescens* (NC_005126); *Serratia marcescens* (NZ_HG326223); *Yersinia pestis* (NC_00314)). Outgroups were selected according to the phylogeny of Gammaproteobacteria by Williams et al. [27]. Gene calling of all genomes was performed using GeneMark version 2.5 [44]. Single copy genes for phylogenetic analysis were retrieved with Orthofinder [45]. Nucleotide and protein alignments for all orthogroups were conducted using MAFFT [46]. Orthogroup alignments that showed evidence of recombination according to the test of Bruen et al. [47] were excluded from further analyses. All remaining protein-as well as nucleotide alignments were concatenated into a supermatrix, resulting into two datasets (available on github XXX). For phylogenetic analyses, the best model for each ortholog partition, as well as the best partition scheme was inferred using IQ-TREE version 1.4.2 [48]. Finally, a Maximum likelihood analyses was conducted for both datasets (proteins and nucleotides) with the same program. Branch support was estimated by using 1000 ultrafast bootstrap replicates [49].

### Endosymbiont host interaction

*Ch. carnea* s. str. develops via three larval instars and a pupal period. Total developmental time from egg to adult was approximately 70 days under our laboratory conditions. Cultivation and all experiments were carried out at 22°C ± 5°C, a photoperiod of 16:8 (light:dark) and 65% ± 5% relative humidity. The cultivation of *Ch. carnea* for the purpose of our experiments started with the second instar larvae obtained from the company Sautter & Stepper GmbH (Ammerbuch-Altingen, Germany). All larvae were reared individually in small round plastic containers (3 cm diameter) and fed with dead moth eggs *(Sitotroga* sp., Katz Biotech, Baruth/Mark, Germany) twice per week. To exclude contaminations from the diet, *Sitotroga* eggs were PCR screened for *Rickettsia* and *Sodalis* infections, which were not detectable in the diet. Adult lacewings were fed with a mixture of honey, water, yeast extract, and sucrose (1:1:1:1) every other day. Adults aged 7 days were then mated by putting 2–4 females and 2–4 males in one cage (38×38×60 cm, total 4 cages). After 5 days females were separated into a plastic container (~ 38 cm^2^), covered with a fine cotton mesh to encourage oviposition and fed every other day with the food mixture described above. The mesh containing eggs was changed every 5 days and stored in a small petri dish (6×1.5 cm). These dishes were checked daily and hatched larvae were collected and reared separately to reduce the rates of cannibalism. After eggs were collected, all mothers were screened via PCR for the presence of *Sodalis* and *Rickettsia* as described above. This allowed us to rear lines of lacewings that were either 1) symbiont free, 2) infected with Rickettsia only, 3) infected with *Sodalis* only or 4) infected with both *Rickettsia* and *Sodalis*. All experiments aimed at comparing these four groups were performed with F2 adults, i.e. after two generations of cultivation in the laboratory.

First, to determine the vertical transmission rate for the endosymbionts *Rickettsia* and *Sodalis* in *Ch. carnea* s. str., 18 *Ch. carnea* s. str. females were investigated (3 with *Rickettsia* only, 2 with *Sodalis* only, 13 with *Rickettsia* and *Sodalis*. For this purpose, all females were mated with *Rickettsia* and y *Sodalis* infected males and reared as described above. After 16 days females were removed and screened for *Rickettsia* and *Sodalis* symbionts as described above. The offspring of those females were collected every day and allowed to develop for 28 days (until stage 2) before being collected and PCR screened for the symbionts. The rate of vertical transmission was determined by calculating number of infected offspring/number of total offspring tested per female.

Second, to test if the symbionts impact reproductive success, 36 female lacewings were investigated (9 with *Rickettsia* only, 8 with *Sodalis* only, 12 with *Rickettsia* and *Sodalis*, 7 without symbionts). *Rickettsia, Sodalis*, and *Rickettsia* and *Sodalis* infected females were mated with males of the same infection status. Only in combined endosymbiont pairs, 4 males were only infested with *Sodalis*. Females without symbionts were mated with *Rickettsia* or *Rickettsia* and *Sodalis* infected males. After a mating period of five days, adults were separated as described above, and females were placed individually in round plastic containers. We counted the number of eggs per female every five days for 45 days and transferred all eggs from one female into a small petri dish. After 45 days, the mothers were screened for symbionts by PCR. Beginning from the first collection of eggs, the hatched larvae were counted visually every day until all eggs were empty or dried out. All larvae were then reared separately. The number of pupae and emerged adults were counted visually every day as well. Finally, using a general linear model with a quasi-Poisson distribution in R [50], we compared the reproductive success for the categories ‘number of eggs’, ‘larvae’, ‘pupae’ and ‘emerged adults’ for the four investigated groups.

Third, to determine, if symbiont titer correlates with reproductive success, we used quantitative real-time PCR (qPCR). To this end, we collected 29 adult females (9 with *Rickettsia* only, 10 with *Sodalis* only, 10 with *Rickettsia* and *Sodalis*). All of them were unmated and of equal age (14 days). Genomic DNA was extracted as described above. A 222bp fragment of *gltA* and 182bp fragment of *groEL* was amplified from *Rickettsia* and *Sodalis*, respectively. Specific primers for these fragments were designed using Prime3 (Table S2, [51]) and their efficiency ensured by creating standard curves. As reference gene to normalize between samples, we amplified a fragment of the single copy nuclear gene *actin*, using primers from Liu et al. [52].

All reactions were run on a PikoReal Real-time PCR System (Thermo Fisher Scientific, Waltham, USA). A 10μl reaction mixture contained SYPR^®^ Green qPCR master mix (2X, Thermo Fisher Scientific, Waltham, USA), 2.5μmol of forward and reverse primer and 10μg DNA. The qPCR program was set as follows: initial incubation at 95°C for 1 min, followed by 40 cycles at 95°C for 15s, 55°C for 15s, and 72°C for 45s.

Differences between groups were determined using an one-way ANOVA with Tukey post-hoc test in R [50]. For statistical analysis, the relative copy number of all genes was normalized by using a log_10_ transformation (Table S4).

## Results

### Endosymbiont screening on population levels in natural and laboratory *Ch. carnea* s. str

To compare distribution patterns of endosymbionts in *Ch. carnea* s. str. in natural and laboratory populations, we screened 148 individuals representing eight natural and three commercially propagated populations. *Rickettsia* infections were found in all natural populations and in total 64% of *Ch. carnea* s. str. individuals were infected (33% to 92% individuals per population infected, Fig. 1). In commercial laboratory populations *Rickettsia* was found in 66% of all screened individuals (25% to 94%). The endosymbiont *Sodalis* was found in 83% (70% to 94%) of all screened individuals in laboratory populations often in combination with *Rickettsia* infections (59%, Fig.1). However, *Sodalis* was not detectable in natural populations.

**Fig. 1.**
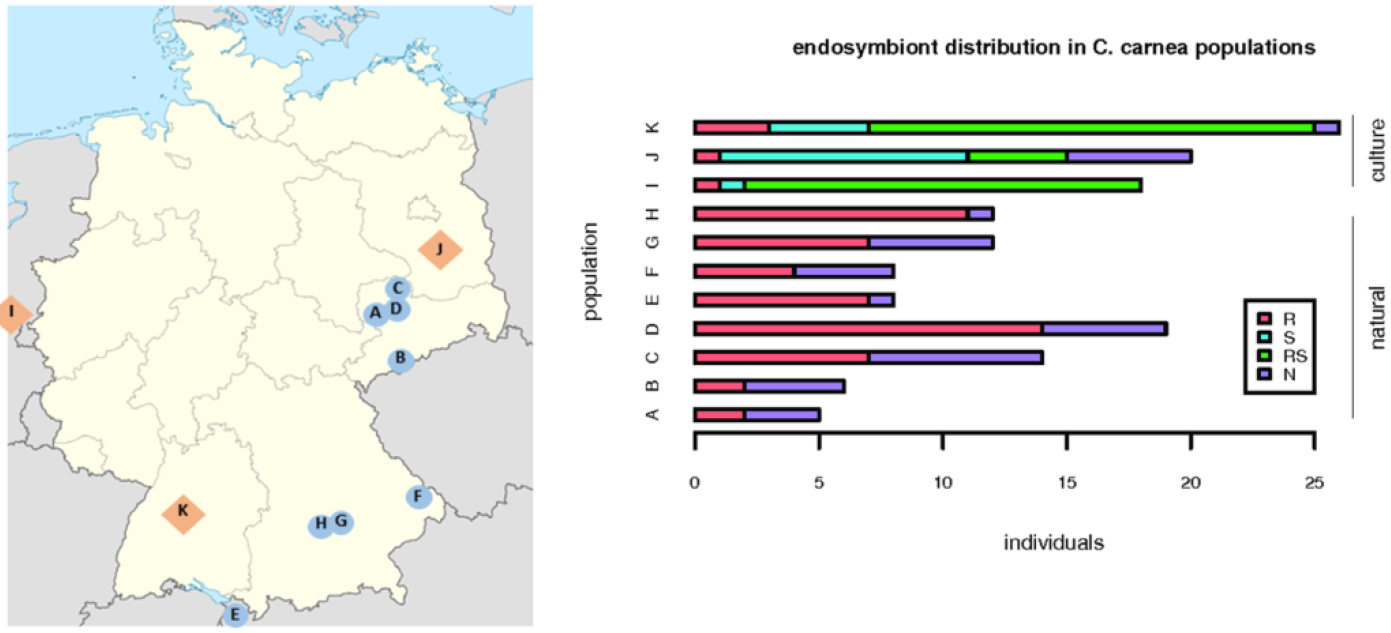
Distribution of *Rickettsia* and *Sodalis* symbionts in *Chrysoperla carnea* s. str. in natural populations (Saxony and Bavaria, Germany, map: circle) and commercial reared populations (map: rhombus). Dark grey: *Rickettsia* infected (R), light grey: *Sodalis* infected (S), black: *Rickettsia* and *Sodalis* infected (RS), white: uninfected (N). Letters in the map highlight founding/rearing places. A: Trages, B: Neudorf, C: Dahlen, D: Püchau, E: Rankweil, F: Schönberg, G: Wippenhausen, H: Kranzberg, I: Biobest, J: Katz Biotech, K: Sautter & Stepper

### Molecular phylogenetic characterization of endosymbionts

A previous phylogenetic analysis demonstrated that the *Rickettsia* strain infecting *Ch. carnea* belongs to the *Rickettsia bellii* clade [26]. For the *Sodalis* strain in *Ch. carnea*, 4,289,304 reads could be used to assemble a draft genome, which was represented by 558 contigs with an N50 of 20,104 and a coverage of ~67x. Based on this draft, the genome of the *Sodalis* endosymbiont is around 4.3 Mbp in size and 5213 coding sequences were identified.

For the phylogenomic analyses, 435 single copy orthologs present in all terminals of our dataset were identified. After removing all ortholog alignments which showed significant signs of recombination, 399 orthologs remained for the final analyses. The concatenated supermatrix consisted of 144,746 amino acid positions. Partitioned Maximum Likelihood analysis recovered a monophyletic group of *Sodalis* strains with a bootstrap support of 100% (Fig. 2). Among *Sodalis*, two reciprocal monophyletic groups were found, both maximally supported. The *Ch. carnea* infecting strain was found as sister taxon of *Sodalis praecaptivus*, in a group that further on contained *Sodalis pierantonius*, *Sodalis* sp. TME1 and *Sodalis* endosymbiont of *Proechinophthirus fluctus*.

**Fig. 2.**
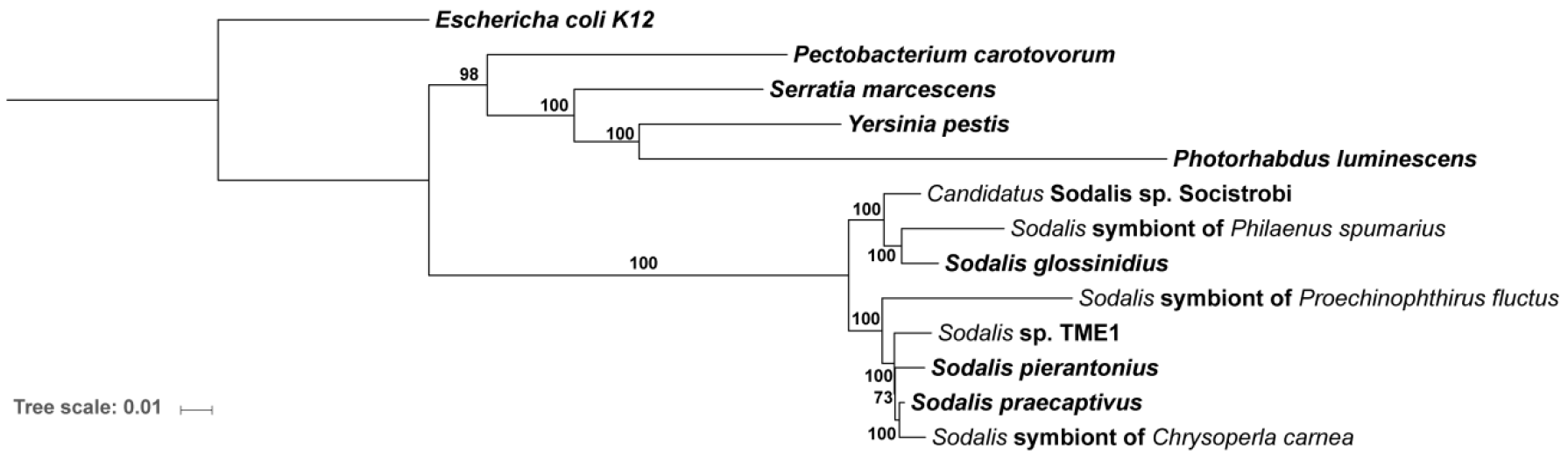
Phylogenomic analyses of 399 proteins alignments single copy orthologs of *Sodalis* strains using Maximum Likelihood as implemented in IQ-TREE. Best models for each gene partition, as well as the best partitions scheme were estimated using IQ-TREE. Bootstrap support from 1000 ultrafast replicates is given at the nodes.

### Endosymbiont host interaction

The rate of vertical transmission estimated from the number of infected offspring divided by the number of total offspring was very high for *Sodalis* (96.3% in single and co-infected lacewings). *Rickettsia* was transmitted at a slightly lower rate (89.0% in single and co-infections, Table 1). In general, the vertical transmission rates were slightly higher in the case of single infections when compared with double infections (93% vs. 87% for *Rickettsia* and 100% vs. 96% for *Sodalis*). Reproductive success differed considerably between groups. We found that the presence of *Sodalis* reduced reproductive output in comparison to uninfected lacewings (Fig. 3, Table 2). The same was true for co-infected lacewings, which showed the lowest reproductive success. No significant differences in performance were found in *Rickettsia* infected *Ch. carnea* compared to uninfected lacewings. In general, the number of hatched larvae was rather low, which might indicate sub-optimal rearing conditions and an effect of this on our results cannot be ruled out (Fig.3, Table S3).

**Table 1.**
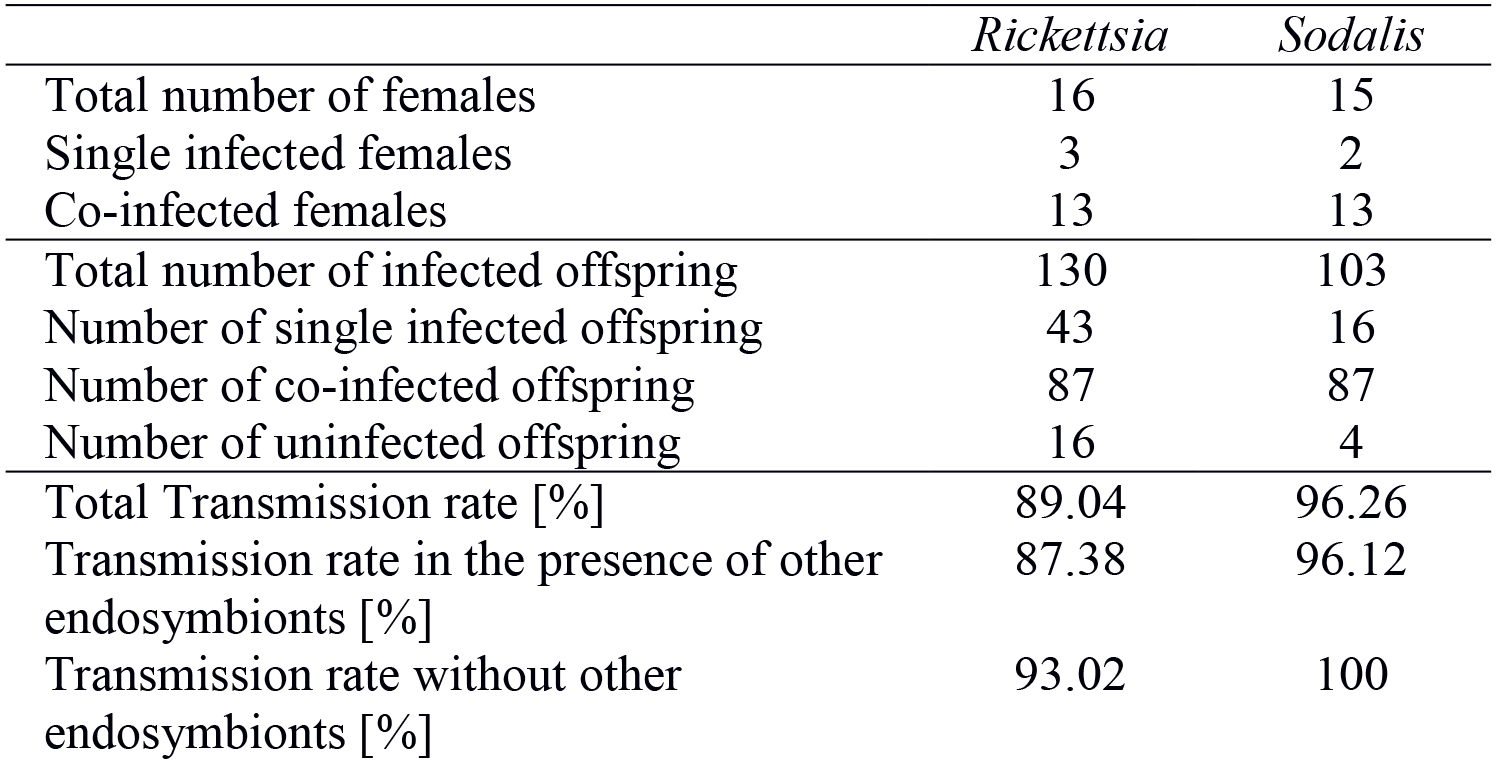
Vertical transmission of single and co-infections of *Rickettsia* and *Sodalis* in *Chrysoperla carnea* s. str.

|  | *Rickettsia* | *Sodalis* |
| --- | --- | --- |
| Total number of females | 16 | 15 |
| Single infected females | 3 | 2 |
| Co-infected females | 13 | 13 |
| Total number of infected offspring | 130 | 103 |
| Number of single infected offspring | 43 | 16 |
| Number of co-infected offspring | 87 | 87 |
| Number of uninfected offspring | 16 | 4 |
| Total Transmission rate [%] | 89.04 | 96.26 |
| Transmission rate in the presence of other endosymbionts [%] | 87.38 | 96.12 |
| Transmission rate without other endosymbionts [%] | 93.02 | 100 |

**Table 2.**
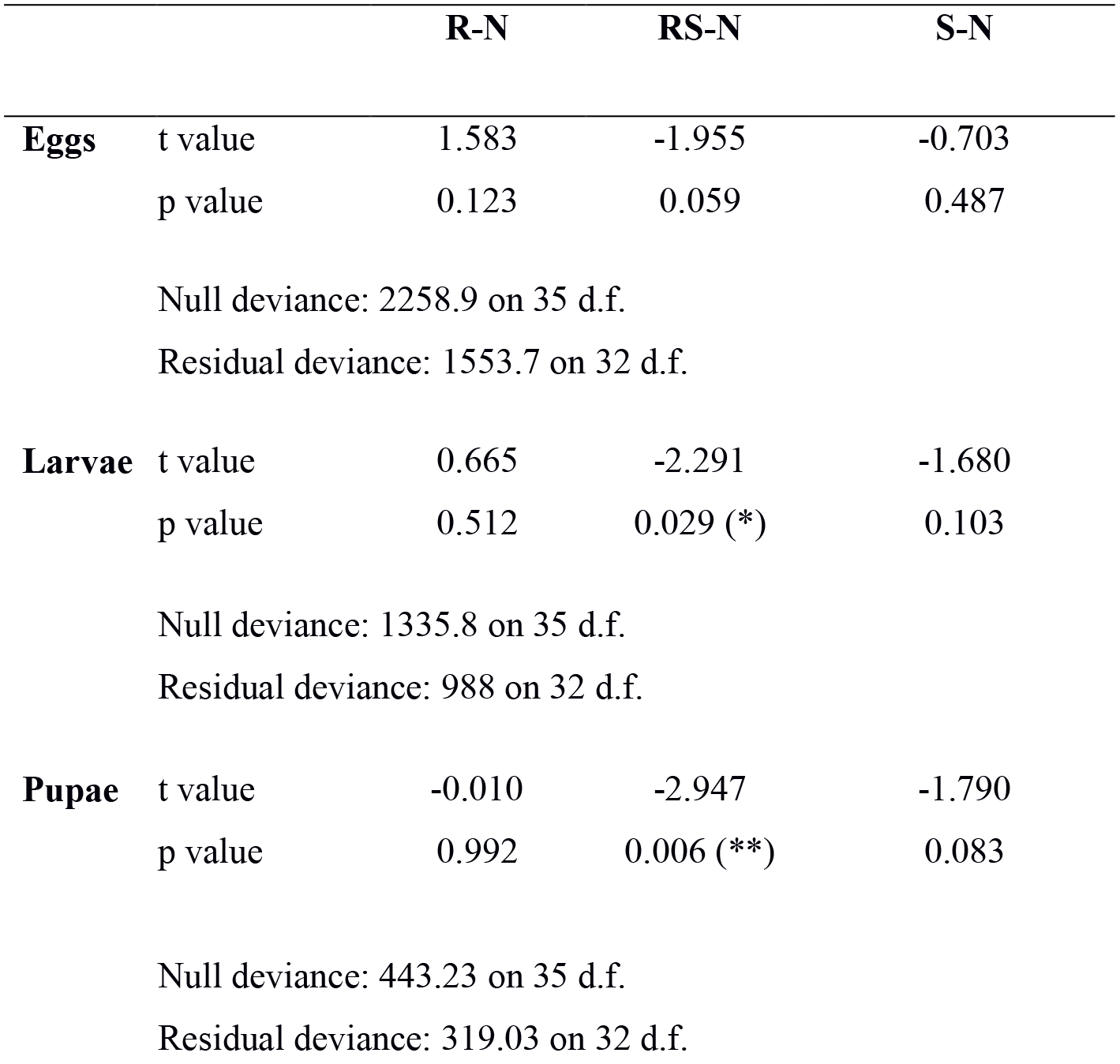
Statistical comparison of laid eggs, viable larvae, number of pupae and emerged adults of *Chrysoperla carnea* s.str. lines of different endosymbiont infections, using a general linear model with a quasi-Poisson distribution. N: no endosymbiont, R: *Rickettsia*, S: *Sodalis*, RS: *Rickettsia* and *Sodalis* infected.

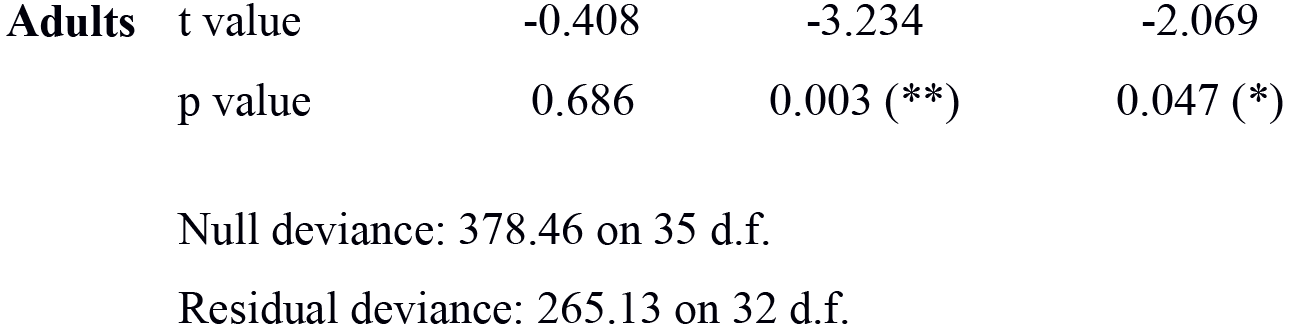

Finally, by using qPCR we found that *Rickettsia* titers were almost uniform across life stages and independent of co-infecting *Sodalis* (Fig. 4). *Sodalis* titers did not significantly differ between single and co-infected adults. However, *Sodalis* titers were significantly reduced in co-infected larvae in comparison to adults (Fig. 4, Table 3). Nevertheless, the highest *Sodalis* titer was observed in adults infected with *Rickettsia*. This line (infested with *Rickettsia* and *Sodalis*) showed the lowest number of laid eggs (Fig. 3). Lacewings with single *Sodalis* infections showed a lower *Sodalis* titers and a higher reproductive success than co-infected lacewings.

**Table 3.**
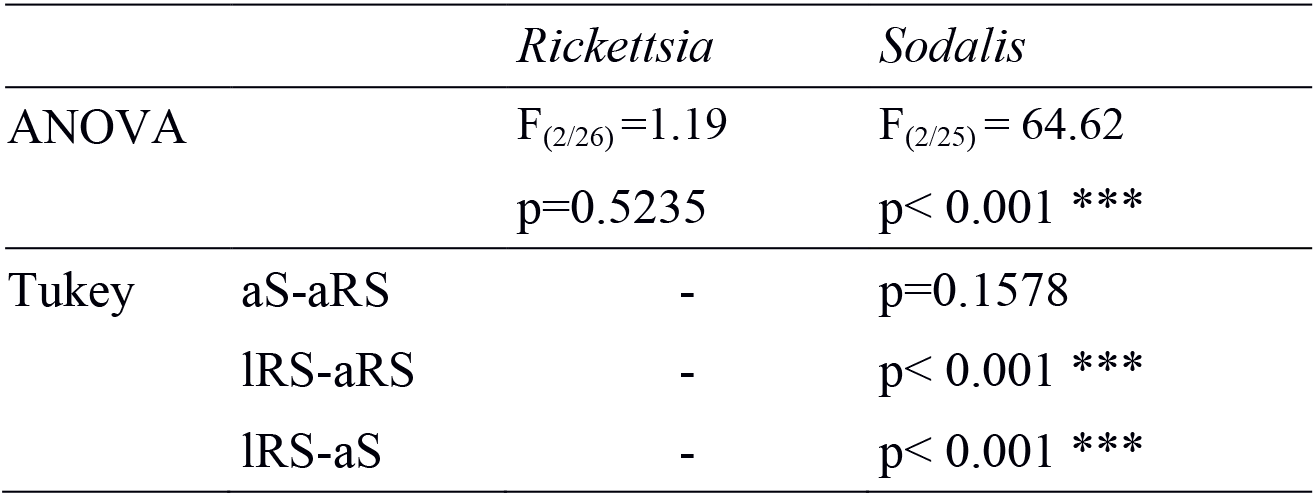
Comparison of copy number of *Rickettsia*, *Sodalis* and co-infected *Chrysoperla carnea* s. str. using a one-way ANOVA and Tukey post-hoc test. a: adult; l: larvae; S: *Sodalis* infection only; R: *Rickettsia* infection only; RS: co-infection with *Rickettsia* and *Sodalis*.

## Discussion

### Population level endosymbiont screening in natural and laboratory *Ch. carnea*

We found two endosymbionts to be common in *Ch. carnea* s. str.: *Rickettsia* and *Sodalis*. In the case of *Sodalis* sp. it is the first record for Neuroptera. However, the most common endosymbiont was *Rickettsia*, which occurred in all tested natural and laboratory population and in all life stages. Our screening of *Rickettsia* infections revealed infection rates ranging from 25% to 94% in both investigated population types (laboratory and natural). *Rickettsia* is a common endosymbiont in arthropods, estimated to be distributed in a quarter of all arthropod species [13]. The infection rate is highly variable in the insect species that were investigated so far. Wild whitefly (*Bemisia tabaci*, Hemimptera: Aleyrodidae) populations showed a *Rickettsia* infection frequency ranging from 22% to 100%, Buprestidae (Coleoptera) 46.3% and the mirid bug species *Nesidiocoris tenuis* (Heteroptera: Miridae) 93% to 100% [53,18,54]. The infection rate we found in lacewings are thus in line with studies mentioned above.

While screening green lacewings for endosymbionts using a metagenomic approach, we also detected *Sodalis*, a well-known gammaproteobacterial endosymbiont of tsetse flies [55], but also detected in other insects, such as stinkbugs, spittle bugs, bird lice, hippoboscid flies, weevils, psyllids or scale insects. [33,32,56]. Interestingly, we detected *Sodalis* in high frequency in *Ch. carnea* s. str. individuals from all studied laboratory populations, but never in natural populations (Fig. 1). A difference in the presence of *Sodalis* in commercially available specimens versus naturally collected specimens was also noticed by Saeed and White in bees [57]. They detected *Sodalis* in only 3 out of 100 individual bees captured in the wild, but in 10 out of 85 individuals when sampling commercially reared. However, differences in endosymbiont infection rate between natural and laboratory populations seem to be host species dependent. Whereas *Wolbachia* infection rates were similar in natural and laboratory vinegar flies *Drosophila melanogaster* (Diptera) populations [58], in tsetse flies, the same *Wolbachia* endosymbiont showed a highly fluctuating infection rate in natural populations and a 100% infection rate in laboratory populations for one tsetse fly species. Another tsetse fly species showed a higher infection rate in laboratory than natural populations [59].

The complete absence of *Sodalis* in natural *Ch. carnea* s. str. populations may be caused by differences in potential selection pressures between laboratory and natural populations, e.g., fluctuating environmental conditions, natural enemies and competition for nutrition with other arthropods. Conceivably, these additional sources of stress are more relaxed or missing under laboratory conditions, which may be favorable for *Sodalis*. In line with this, our data suggest that *Sodalis* cause fitness costs for *Ch. carnea* s. str., and it could be assumed that this so far uncharacterized impact to be less severe in the laboratory. Based on the fitness costs, *Sodalis* may be faster eliminated in natural populations than in laboratory populations, where inbreeding and stable conditions may enhance the transmission rate of endosymbionts. E.g., a correlation of temperature with the rate of transmission has been reported for several bacteria [60,61]. Higher *Sodalis* infection frequency was detected in weevils living at localities of higher temperature than of lower temperature [62]. Also a lower mortality in the presence of *Sodalis* in laboratory cultures or reinfections from the environment are conceivable. However, given the current state of knowledge it can only be speculated why *Sodalis* is only present in laboratory populations of the green lacewings.

### Molecular phylogenetic characterization of endosymbionts

Endosymbionts show a broad range of interactions with their hosts, including obligate or facultative mutualism or parasitism [63]. To understand more about the impact of the endosymbionts *Rickettsia* and *Sodalis* on their *Ch. carnea* host, both endosymbionts were characterized genetically. *Rickettsia* occurs in many diverse arthropod orders and it is subdivided into 13 lineages [16]. We found the same strain (*R. bellii* group) in the here investigated green lacewings as Gerth et al. [26] already reported for different populations the same species. Interestingly, diverse *Rickettsia* lineages infect Neuroptera and they are distributed in species-specific manner [26].

To characterize the *Sodalis* endosymbiont, we used a phylogenomic approach. This analysis showed that this strain is closely related with *Sodalis praecaptivus* [64]. Several other close relatives of this species have already been identified in different insect hosts [65,66]. Whereas basically all other known *Sodalis* strains have been described as primary or secondary endosymbionts of insects, *S. praecaptivus* was isolated from a human wound which was the result from an accident with a tree branch. The *S. praecaptivus* strain is regarded as a free-living member of *Sodalis* and it has been shown that its genome is with 5.17 mbp the largest of all so far sequenced *Sodalis* strains. Intriguingly, the genome size of other *Sodalis* strains seem to correlate with the dependency to its host. The *Sodalis*-like primary endosymbiont of the spittlebug *Philaenus spumarius* has with 1.39 mbp the smallest genome of all known strains [56,67]. With 4.3 mbp, the draft genome of the green lacewing *Sodalis* strain is comparable in size to those of Candidatus *S. pierantonius* (4.5 mbp), a secondary endosymbiont of the rice weevil [68], and *S. glossinidius* (4.2 mbp), a secondary endosymbiont of the tsetse fly [69]. It has been hypothesized that *Sodalis* strains adapted independently to an endosymbiotic life-style with different insect hosts, resulting in a reduction of genome size and complexity [68,70]. However, a more contiguous assembly of the green lacewing *Sodalis* strain is necessary for a detailed analysis of the state of its “genome degeneration”.

### Endosymbiont host interaction

Based on the high infection rate in several *Ch. carnea* s. str. populations, the vertical transmission rate of both endosymbionts were investigated. *Rickettsia* and *Sodalis* showed a high rate of vertical transmission (89–96%, Table 1). The *Rickettsia* transmission rate is consistent with an earlier study in whiteflies under laboratory conditions [21]. Slightly lower rates were found in studies of tsetse flies (*Glossina morsitans*) for *Sodalis*, (67–75%, [71,35]). In our study, single infections were transmitted at a slightly higher rate than double infections. By sharing the same host, endosymbionts have to compete for nutrients and space either by sharing resources or evolving niches, e.g. inhabit special cells or organs [72,73]. This phenomenon was found in tsetse flies, where *Wolbachia* only infects oocytes, *Wigglesworthia* bacteriocytes and milk gland, and *Sodalis* several organs [74].

Negative fitness impacts are a prevalent phenomenon associated with endosymbionts [12]. We therefore investigated if *Rickettsia* and/or *Sodalis* impact host reproductive success in *Ch. carnea*. In the present study we found no significant impact for single *Rickettsia* infections on the reproductive success of its host (Fig. 3). However, a trend towards an increase in the number of laid eggs is visible when compared to uninfected lacewings. *Rickettsia* manipulate other insects in a negative or positive way. It has a negative impact in aphids on body weight, fecundity and longevity [75,76] and a positive impact in whiteflies and leeches on body size, number of offspring, development and survival rate [77,21]. In the present study *Sodalis* seems to have a detrimental effect on number of viable offspring in *Ch. carnea*. This impact on fecundity and pupal emergence rate was not found in tsetse flies. In those hosts, S*odalis* establishes trypanosome infections and longevity [78,35,79]. The impact of *Sodalis* on other insects is less well understood. However, the low larval hatching rate in our study indicates sub-optimal rearing conditions and an effect of this on our results cannot be ruled out.

**Fig. 3.**
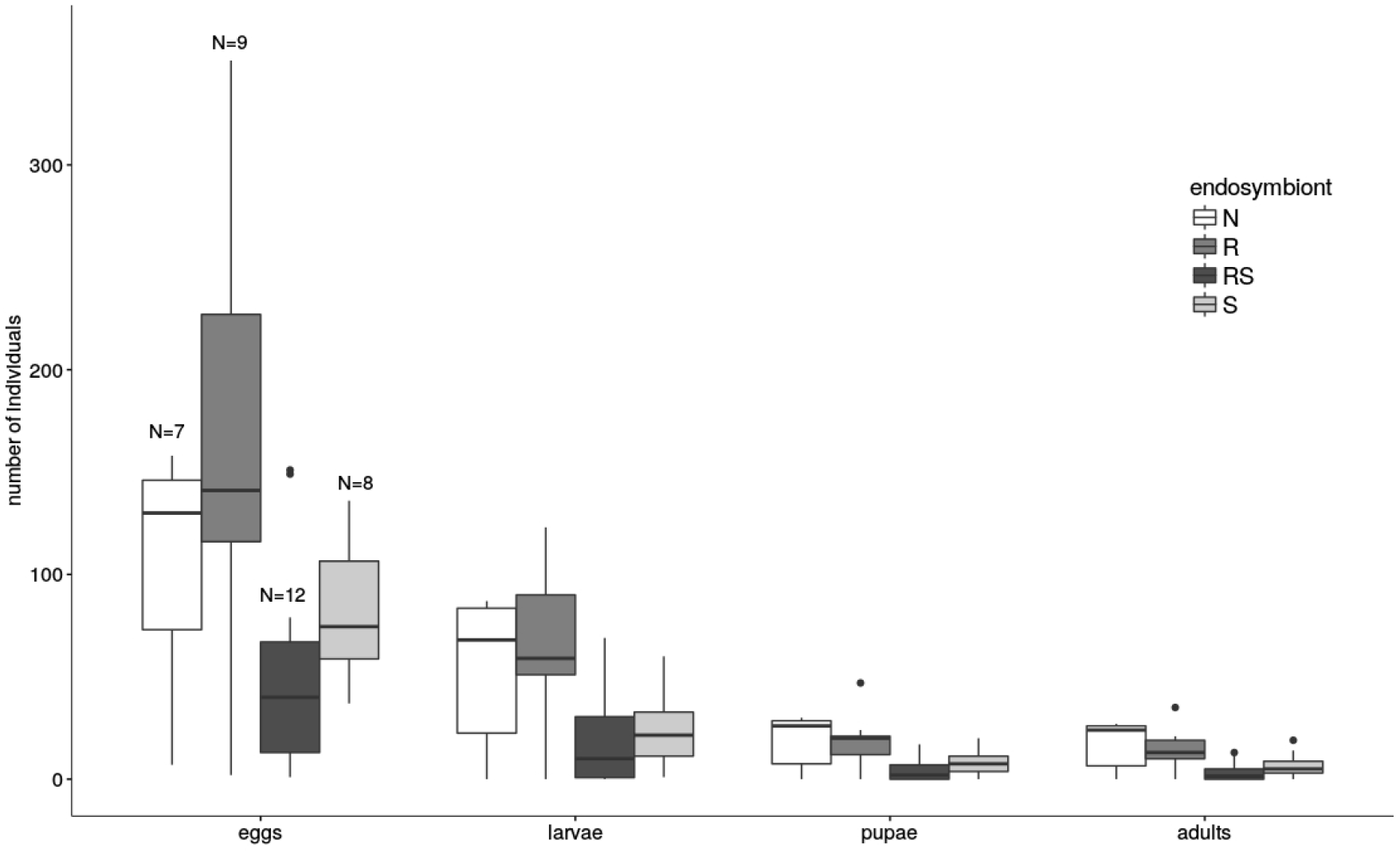
Number of laid eggs, viable larvae, pupae and adults of *Chrysoperla carnea* s. str. of lines differing in levels of endosymbiont infections. White: non-infected, dark grey: *Rickettsia*, light grey: *Sodalis* or black: co-infected with *Rickettsia* and *Sodalis*.

The co-occurrence of *Rickettsia* and *Sodalis* found in *Ch. carnea* s. str. was also reported from weevils and lice [62,80]. In our study, co-infections showed a detrimental effect on the reproductive success, partially stronger than in single *Sodalis* infections. To test if there is a connection between endosymbiont density and reproductive success, *Rickettsia* and *Sodalis* titers were measured in single and co-infected lines. *Rickettsia* showed constant titers independent of the infection type (single-or co-infected, Fig. 4). Based on similar density in larval and adult lacewings, we presume a constant *Rickettsia* infection density across life stages. Interestingly, single *Sodalis* infections showed a tendency to lower densities than co-infections (Fig. 4). *Sodalis* densities in adult lacewings correlated negatively with the number of laid eggs. Co-infected lacewings showed the highest *Sodalis* titer and laid the least eggs. We suggest that *Sodalis* is causing a fitness disadvantage of adult lacewing host and this negative effect on reproductive success increases with a higher *Sodalis* density. It is conceivable that endosymbionts compete for resources and space by sharing the same host in coinfections [72,73]. However, *Sodalis* infections were not found in all host life cycle stages in other insects. For example in cereal weevils (*Sitophilus*, Coleoptera) the endosymbiont is involved in the cuticle synthesis in young adults, while afterwards the endosymbiont is eliminated [31]. In contrast, we found that co-infected larvae showed a significantly lower *Sodalis* density compared to adults. One hypothesis is that in *Ch. carnea* s. str. *Sodalis* mainly occur in reproductive organs such as ovaries and gonads, as also reported for stinkbugs [29]. In *Ch. carnea* s. str. larvae the *Sodalis* density can be reduced due to the fact that these organs are not developed until this stage. However, this preliminary result needs further investigation, especially regarding the relevance of the symbionts and their function in several life circle stages.

**Fig. 4.**
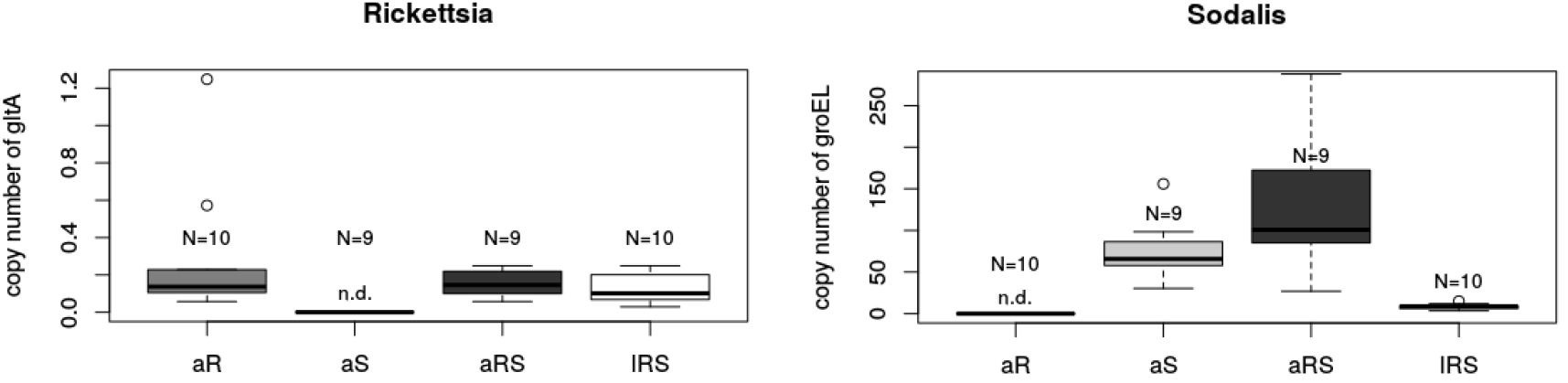
Copy number of *Rickettsia* specific *gltA* (left) and *Sodalis* specific *groEL* genes (right) in *Rickettsia* or *Sodalis* single or co-infected *Chrysoperla carnea* s. str. a: adult; l: larvae; S: *Sodalis* infection only; R: *Rickettsia* infection only; RS: co-infection with *Rickettsia* and *Sodalis*.

This work is a first step in studying the distribution and fitness impact of endosymbionts in the common green lacewing *Ch. carnea* s. str., a species frequently used in biological pest control. The negative fitness effect found in this study may have an important impact on commercial rearing and it should be explored if treating *Ch. carnea* s. str. with antibiotics may improve the rearing success and efficiency. However, *Ch. carnea* s. str. is not dissociable from its microbiome and may be strongly influenced by its symbionts and symbiont-symbiont-interactions, especially symbiont-symbiont interaction are rarely understood.

## Acknowledgments

We thank Katharina Grosser for her support in cultivation, Christian Ristok for helpful comments on statistics and Anne Weigert for supporting the Illumina sequencing. Felix Wäckers of BioBest is thanked for providing *Ch. carnea*.

